# fastMSA: Accelerating Multiple Sequence Alignment with Dense Retrieval on Protein Language

**DOI:** 10.1101/2021.12.20.473431

**Authors:** Liang Hong, Siqi Sun, Liangzhen Zheng, Qingxiong Tan, Yu Li

## Abstract

Evolutionarily related sequences provide information for the protein structure and function. Multiple sequence alignment, which includes homolog searching from large databases and sequence alignment, is efficient to dig out the information and assist protein structure and function prediction, whose efficiency has been proved by AlphaFold. Despite the existing tools for multiple sequence alignment, searching homologs from the entire UniProt is still time-consuming. Considering the success of AlphaFold, foreseeably, large-scale multiple sequence alignments against massive databases will be a trend in the field. It is very desirable to accelerate this step. Here, we propose a novel method, fastMSA, to improve the speed significantly. Our idea is orthogonal to all the previous accelerating methods. Taking advantage of the protein language model based on BERT, we propose a novel dual encoder architecture that can embed the protein sequences into a low-dimension space and filter the unrelated sequences efficiently before running BLAST. Extensive experimental results suggest that we can recall most of the homologs with a 34-fold speed-up. Moreover, our method is compatible with the downstream tasks, such as structure prediction using AlphaFold. Using multiple sequence alignments generated from our method, we have little performance compromise on the protein structure prediction with much less running time. fastMSA will effectively assist protein sequence, structure, and function analysis based on homologs and multiple sequence alignment.

## 1 Introduction

The large volume of available biological data in recent years have promoted computational biological research, such as protein structure and function predictions, protein secondary structure analysis, unknown protein sequence discovery, and phylogenetic reconstruction [29, 46, 55, 53, 9, 3]. As an essential and significant computational procedure for these analysis tasks, Multiple Sequence Alignment (MSA) refers to identifying and aligning the evolutionarily related biological sequences, which usually need to take evolutionary events (such as deletions, insertions, rearrangements, and mutations) into consideration [10]. It has become an indispensable step for biological data analysis, whose quality severely influences the performances of many biological analysis tasks [56, 28, 17, 30, 3].

MSA originates from the pairwise sequence alignment. To solve the pairwise problem, we will find an alignment that maximizes the similarity between two sequences [16]. One typical way of estimating similarity is to summarize substitution matrix scores of each aligned residue pair, which can be exactly solved from a mathematical perspective [16]. However, MSA is much more complex, which needs to find an alignment that maximizes the sum of similarities for all sequence pairs. In fact, finding an optimal alignment for multiple sequences from the mathematical view is a complex optimization (NP-complete) problem because it needs to identify the best MSA from the entire set of possible alignments [12]. Hence, heuristic methods are usually needed. The most popular traditional MSA method is a progressive alignment algorithm, which generates a phyletic tree and aligns a set of sequences [19, 10]. This method uses a phyletic tree to align multiple sequences while the obtained alignments are then applied to adjust the tree. Along this line, some methods are designed to construct global pairwise alignment and introduce tree-based progressive strategies, including ProbCons [14], T-Coffee [33], and ClustalW [42]. To improve quality, the iterative strategy is further introduced to repeatedly adjust the tree and alignment process until both of them converge [45], such as Clustal Omega [40], MUSCLE [15], and MAFFT [24]. These designs promote the alignment quality of multiple sequences to some extends. However, these methods need to optimize an alignment by considering a set of pre-defined constraints, which is an NP-complete problem and is nearly impossible when the number of sequences is large [10]. Thus, these methods are slow and can only handle datasets with small instances [12, 25].

In recent years, some new MSA algorithms are proposed by considering the structure information of already existing protein sequences [12, 10], such as 3DCoffee [34], MICAlign [51]. Their basic idea is that the structures of protein sequences usually evolve much more slowly than sequences themselves [26]. Hence, they believe that if a model can consider the structure of sequences, this model has great potential of achieving better results than other models that only rely on sequence alignment [26, 12, 11]. Armougom et al. introduce a server termed EXPRESSO to automatically select templates by running BLAST [2, 8] to identify close homologous by considering the structural information of sequences in the PDB database [4]. The incorporation of structure information can help promote the alignment effeteness of multiple sequences. However, these algorithms rely heavily on the structure information obtained from known protein sequences, which makes them have weak scalability because the structure of many protein sequences may not be available. What’s worse, these methods often have low speed, which will further limit their real-world applications, especially for large protein databases.

To handle protein databases with a huge number of sequences [7, 41], people are developing faster algorithms to effectively decrease the running time of models. Buchfink et al. introduce the diamond algorithm that is built upon the double indexing for protein sequence alignment, which achieves a similar degree of sensitivity with the gold standard BLASTX but is much faster [7]. Huson et al. present a new method named protein-alignment-guided assembly for gene-centric assembly, which provides fast access to each gene family in MEGAN [20]. Steinegger et al. design a single-instruction multiple-data vectorized implementation version of Viterbi algorithm to align protein sequences, which accelerates the search methods HHsearch and HHblits [41]. However, These methods are often designed for relatively small databases, which can not meet the needs of dealing with a very large database. For example, the recently proposed AlphaFold2 needs to obtain the MSA for every protein in the UniProt database, which contains 250 million sequences and provides very good chances to improve the alignment quality [22, 44, 6]. And foreseeably, obtaining the MSA for a huge number of sequences using the large database will be very common in the post-AlphaFold era. However, using traditional methods is too slow to perform such searching. Therefore, it is still urgent to develop an algorithm that is able to efficiently complete the searching procedure for very large databases and thereafter perform the MSA tasks in a relatively short time.

To deal with the above issues, we propose a novel method, which is orthogonal to all the previous methods. This fastMSA framework, consisting of query sequence encoder and context sequences encoder, can improve the scalability and speed of multiple sequence alignment significantly. Firstly, to prepare the training data for the proposed method, we utilize the query sequences to search from UniRef90 using JackHMMER v3.3 [50] and build the resulted MSAs as ground truth. Secondly, in the encoder part, we design a transformer-based query sequence and context sequence encoder with learnable parameters. It is designed to learn the sequence embedding, where the contrastive learning strategy is introduced to significantly boost the training efficiency. Thirdly, during the inference time, to boost the speed of the proposed algorithm for conducting the large-scale searching, we apply the context encoder to encode all the sequences to fixed-length vectors to search for top relevant targets. Then, we compare the similarity scores between the encoded query and all the sequences in the database, which are utilized to rank the sequences and retrieve the top-*K* most similar sequences. We will perform the standard homolog searching and MSA on the top-*K* most similar sequences. Extensive experiments suggest that we can recall around 70% homologs with only top-200k sequences, which reduces the database size by 350 folds. It can lead to around 34 folds speed-up. Moreover, our method is compatible with the downstream tasks, such as structure prediction using AlphaFold. Using multiple sequence alignments generated from our method, we have little performance compromise on the protein structure prediction with much less running time. The proposed tool, fastMSA, will effectively assist protein sequence, structure, and function analysis based on homologs and multiple sequence alignment.

## 2 Related Work

### 2.1 Multiple Sequence Alignment

Multiple Sequence Alignment (MSA) has become a hot research area a long time ago [43, 16]. Since the alignment of multiple sequences needs to optimize the sums-of-pairs evaluation schemes, this is a NP-complete problem, where heuristics methods are often required [10]. Based on this consideration, the most common one is the progressive alignment algorithm initially proposed in 1984 [19], which incorporates all the input sequences into the final model sequentially by following an inclusion order that is pre-defined in a guide tree. This method needs to conduct a pairwise alignment for two sequences, two profiles, or a sequence and a profile, at each node, which is often tackled via the global dynamic programming alignment algorithm [32]. After this, more similar algorithms that combine a global pairwise alignment method and a tree-based progressive strategy are designed, such as ProbCons [14], T-Coffee [33], and ClustalW [42].

To improve the prediction performance, the iterative strategies are further introduced to dynamically estimate both the tree and alignment components by repeatedly using dynamic programming until both components converge [45]. Based on this strategy, many algorithms are proposed for the alignment of multiple sequences, including Clustal Omega [40], MUSCLE [15], and MAFFT [24]. At the same time, some methods focus on improving the guide tree of MSA algorithms, which determines the order in which these sequences should be incorporated [10]. Among these methods, Neighbor Joining (NJ) [38] and Unweighted Pair Group Method with Arithmetic Mean (UPGMA) [31] are most widely used, which improve the performance for the alignment of sequences to some extent. However, these methods need to optimize an alignment by considering a set of pre-defined constraints, which is a Maximum Weight Trace problem and is NP-complete [10]. As a result, these methods can only deal with protein sequences datasets with small instances, such as only a few hundred sequences [12, 25]. Therefore, these methods are not able to meet the need of analyzing thousands or even millions of protein sequences in the new data era.

Another way to perform multiple sequence alignments is taking the structure information of already existing protein sequences into consideration. These methods are termed as structure-based MSA [12, 10], such as 3DCoffee [34], MICAlign [51], and EXPRESSO [4]. Because the structures of protein sequences often evolve more slowly than sequences themselves [26], structure-based MSA algorithms are usually able to achieve better performance than methods that only consider sequence alignment [12]. By learning the structure-based multiple sequence alignment from public datasets, Matthieu et al. can successfully identify clear patterns in correlation of crucial residues [11]. However, these methods rely heavily on the structure information of known protein sequences, which may not always be available especially when an increasing amount of new protein sequences are discovered every day.

### 2.2 Unsupervised Protein Language Model

Recent advances in large-scale language model pre-training, such as BERT [13] or GPT [35], have brought various breakthroughs in natural language processing. In order to train the model with minimal human labeling efforts, the training data is constructed by corrupting the sentences, i.e., replacing the original words with a special token, then the task simply is to reconstruct the sentence from the corrupted one.

Similar ideas are then adapted into protein sequences and have emerged as a promising approach in the field. To transfer knowledge between structurally related proteins, Bepler & Berge [5] proposed to use a bidirectional LSTM model and multi-task framework to encode the structural information. A transformer-based language model [36, 37], ESM-1b, is introduced by Rao et al., and demonstrates that information learned from protein sequences alone could greatly benefit for numerous downstream tasks, such as secondary structure prediction, contact prediction and etc, which surpasses the performance of LSTM-based [1, 18] language models by a large margin. Rao et al. showed that MSA transformer could capture more information by integrating MSA into the model, and achieved state-of-the-art results on multiple benchmarks. In addition, the same language model objective is also applied in AlphaFold 2 [22], and helps to further boost the accuracy of structure prediction.

Recently, dense retrieval based on language model in text domain has been explored and achieved state-of-the-art performance in various tasks, such as open-domain question answering and dialog response generation [23, 52, 27, 54]. Karpukhin et al. [23] first proposed to use a bi-encoder architecture to encode query and document respectively, then retrieve document by a similarity score based on dot product. ANCE [52] proposes to use hard negatives to further improve the performance on retrieval. The architecture was later adapted into text generation domain [54, 27], and became a core component for the systems to alleviate the hallucinated facts problem.

## 3 Methods

In this section, we present the proposed fastMSA framework, which consists of two transformer-based models as query sequence encoder and candidate sequences (sequences from a very large database) encoder, as shown in Fig. 1. We first introduce how to construct training data from raw sequences, then present the objective function and detailed inference pipeline, as shown in Fig. 2.

**Figure 1:**
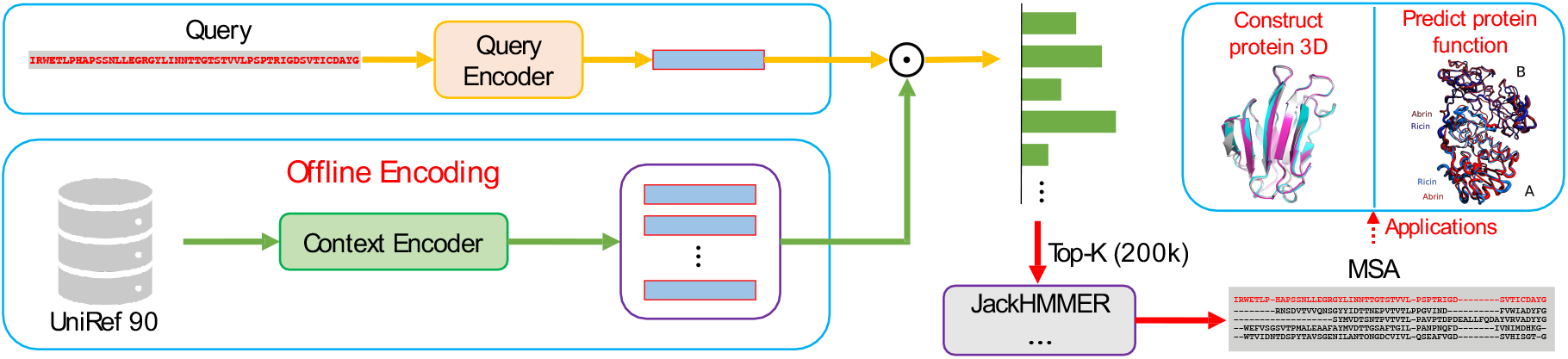
Overview of the inference pipeline for fastMSA. The top-*K* most similar sequences will be retrieved using dot product, then JackHMMER is applied on this small retrieved dataset to build the MSA for further tasks, such as 3D structure prediction or protein function prediction. Before retrieval, the UniRef90 can be encoded into vectors offline, and it will NOT affect the inference time of building MSA.

**Figure 2:**
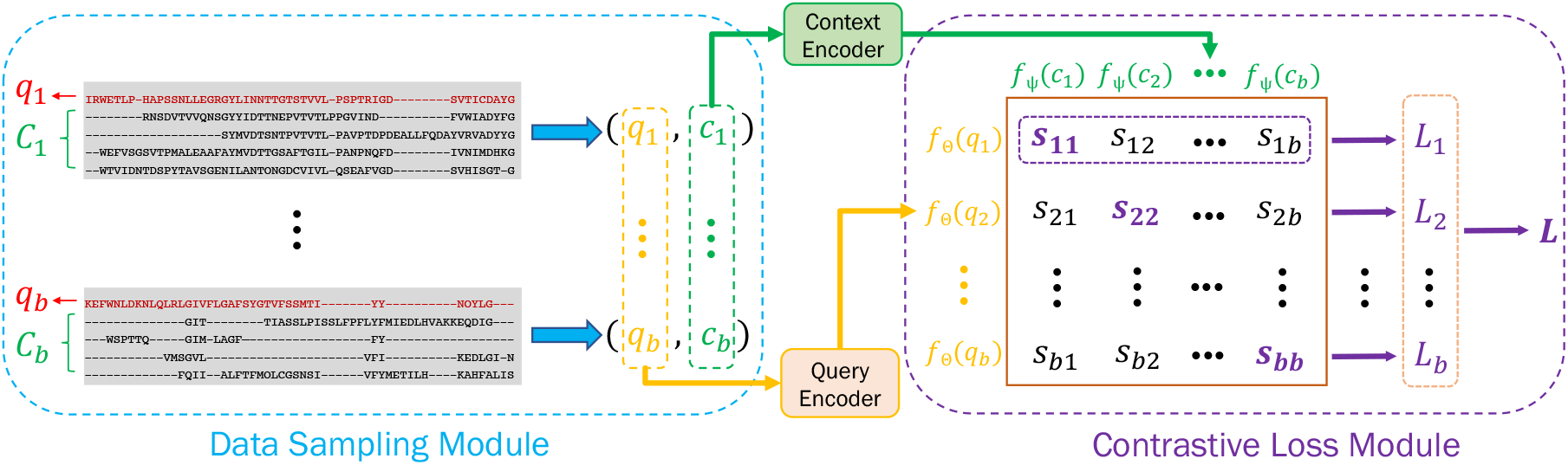
Training pipeline for the proposed method. In the Data Sampling Module, a batch of positive pairs, (*q*_1_, *c*_1_), …, (*q*_*b*_, *c*_*b*_), are sampled from MSAs. Then, a similarity matrix *S* is computed using the contrastive loss, in which each element *s*_*ij*_ = *f*_Θ_(*q*_*i*_)^*t*^*f*_Ψ_(*c*_*j*_). Intuitively, the diagonal of the similarity matrix is trained to be larger than the corresponding off-diagonal elements in the same row since they represent all of the positive pairs.

### 3.1 Pre-training Dataset Construction

To build the training dataset for the proposed method, we first used the query sequence (denoted as *q*) to search from UniRef90 using JackHMMER v3.3 and build the MSA as ground truth. Denote the resulted MSA as 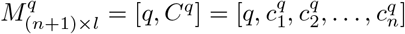, where *l* is the sequence length; the first sequence, *q*, is the query, and the rest *n* sequences, 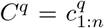, are searched homologs, 1 : *n* denotes set {1, 2, …, *n*}. For simplicity, we drop the superscript *q* if context is clear. Since *q* and *c*_1:*n*_ are from the same MSA, (*q, c*) are treated as a **positive** pair for any *c ∈* {*c*_1_, …, *c*_*n*_}. Note that the amount of training data could be enormous considering the data construction process only depends on JackHMMER to build the MSA.

### 3.2 Bi-Encoder Model and Contrastive Learning

We denote the transformer-based query sequence and candidate encoder as *f*_Θ_ and *f*_Ψ_, respectively, where Θ and Ψ are learn-able model parameters. Given a paired data, say (*q, c*) from the previous section, the query encoder *f*_Θ_ maps *q* into a *d*-dimensional vector, *f*_Θ_(*q*) = *h*_*q*_ ∈ ℝ^*d*^; and the candidate encoder maps *c* into a vector with same dimension, i.e., *f*_Ψ_(*c*) = *z*_*c*_ ∈ ℝ^*d*^. In fact, the transformer encoder will encode the sequence into a series of vectors with the same length, and we simply choose the first vector as its sequence representation, and rely on the training procedure to learn the sequence embedding. Finally, the inner product between these sequence embeddings,

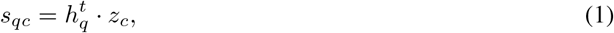

is computed as the similarity score between query *q* and candidate *c*. In addition, both query encoder and candidate encoder are initialized with ESM-1b since the model is already well pre-trained with billions of protein sequences.

Following contrastive learning approaches, we further propose to use in-batch data as negatives, which could significantly boost the training efficiency compared to actual sampling for negatives. More specifically, for a batch of paired data (*q*_1_, *c*_1_), …, (*q*_*b*_, *c*_*b*_), where *b* is the batch size, the negative samples for query *q*_*i*_ are **all** other candidates within the same batch, i.e., all *c*_*j*_ where *j* ≠ *i*. Please refer to the Data Sampling Module in Fig. 2 for more details. Suppose the query and candidate encoder map these queries and their corresponding candidates into matrices, [*h*_1_, *h*_2_, …, *h*_*b*_] = [*f*_Θ_(*q*_1_), *f*_Θ_(*q*_2_), …, *f*_Θ_(*q*_*b*_)] ∈ ℝ^*b*×*d*^, [*z*_1_, *z*_2_, …, *z*_*b*_] = [*f*_Ψ_(*c*_1_), *f*_Ψ_(*c*_2_), …, *f*_Ψ_(*c*_*b*_)] ∈ℝ^*b*×*d*^, the similarity matrix between all queries and homologs therefore are

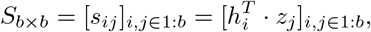

which can be computed efficiently by a simple matrix multiplication. Then, for each query, say for *q*_*i*_, the task is to identify the homolog *c*_*i*_ among all the other negatives candidates *c*_*j*_, *j ≠ i*, which is a *b*-way classification problem. The objective function for query *q*_*i*_ therefore can computed as

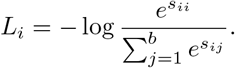

And the final objective function for the batch is the average of all queries, which is 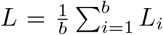. Please refer to the Contrastive Loss Module in Fig. 2 for an illustration.

### 3.3 Online Inference

During the inference stage, we first apply the candidate encoder to encode all the sequences from UniRef90 into vectors and save them into memory. Though it is time-consuming to encode millions of sequences, this encoding step only needs to be done once and can be finished offline. Then, similarity scores between the encoded query and all the sequences in the database are computed based on the equation (1). Finally, the sequences are ranked by the similarity score, and top-*K* most similar sequences will be retrieved for the next stage. Instead of computing the score between the query and all the sequences in the database, this optimization problem can be efficiently solved by approximation packages, such as FAISS [21]. It is worth mentioning that to use the fast search algorithm in FAISS, the similarity score in the equation (1) has to be decomposable, which is the underlying reason why we use dot product as the metric other than a more complex neural network. The retrieved sequences will be used as inputs to JackHMMER to do the final MSA for the input query. Please refer to Fig. 1 for more details about the whole pipeline.

## 4 Results

### 4.1 Dataset Preparation and Hyper-parameters

Because of the limitation in the availability of computation resources, we only managed to generate MSAs on a small dataset, namely CATH [39], which contains around 33k sequences. JackHMMER was utilized to iteratively search for candidate sequences in UniRef90 and align these candidate sequences to the MSA. Then, we constructed our sub-set of UniRef90 by combining all of the top-1k similar sub-sequences in each MSA and referring them back to the original sequence. After these procedures, the obtained sub-set covers 11M distinguished sequences in total.

JackHMMER is then adopted to deal with the obtained distinct sequences. In our implementation, the evaluation threshold and the number of iterations of JackHMMER are set as 10^−3^ and 3, respectively, while other hyper-parameters are set as the default values. In the training process, the model is trained for 100 epochs, and the warm-ups strategy is adopted by linearly increasing the learning rate from 10^−6^ to 10^−4^ gradually and then decaying the rate at 30 and 60 epochs multiplying by 0.1. We use 4 NVidia V100 GPUs, each of which hosts 64 paired data. Cross-batch negative is used to expand the actual batch size to 256.

Three test sequence datasets, namely CASP13, CASP14, and CAMEO, are introduced to evaluate the performance of our trained model, please refer to Table 1 for more statistics of the data. To be specific, the domain level sequences are collected from the CASP13 and CASP14 competitions for MSA retrieval and 3D structure construction. The CAMEO dataset contains all the protein sequences in a half-year period (from 2020-09-07 to 2021-05-01) from CAMEO online 3D structure prediction evaluation.

**Table 1:** The overview of three test datasets, CASP13, CASP14, CAMEO and training set CATH.

| Dataset | #sequences | min length | max length | average length |
| --- | --- | --- | --- | --- |
| **CASP13** | 122 | 31 | 842 | 178.12 |
| **CASP14** | 102 | 35 | 838 | 193.80 |
| **CAMEO** | 297 | 31 | 1176 | 280.13 |
| **CATH** | 32658 | 9 | 1942 | 148.23 |

### 4.2 Retrieval Recall Rate

To evaluate the performance of the proposed fastMSA method, we calculate the recall at top 200k, 400k, 1M, 2M for CASP13, CASP14, and CAMEO, respectively. Corresponding retrieval results are presented in Fig. 3a, which clearly indicates that the proposed fastMSA can effectively decrease the time taken to build MSAs using JackHMMER. It is worth noticing that when computing the recall, all the sequences that appear in the same MSAs and the number of each greater than 10 are regarded as ground truth. We discard the rest queries because the results will highly fluctuate otherwise.

**Figure 3:**
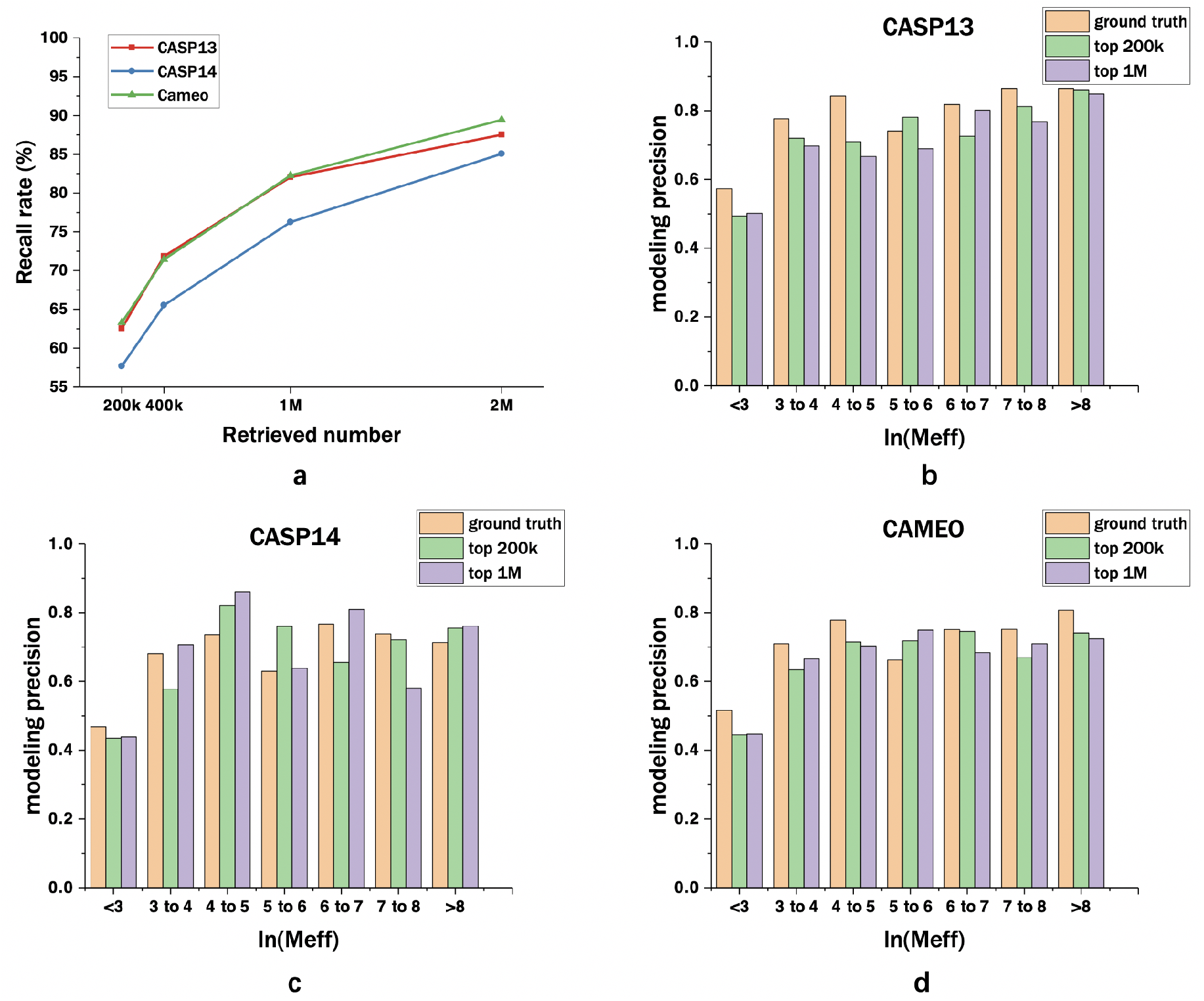
**a**: The recall rate on four data sets. We evaluate our method on three data sets with four different retrieved number: top 200k, 400k, 1M and 2M. **b-d**: Different modeling precision with respect to binned Meff. We compare the ground truth with our method. *>* 9 ground truth column is left blank because ground truth with ln(Meff)*>* 9 does not exist.

Then we analyze the effectiveness of the proposed method for decreasing the time needed for building MSAs. Note that the whole dataset contains around 11M sequences, by reducing the number of sequences by 55 times (11M to 200k), our method retrieves at least 55% of ground truth sequences for CASP 14, and at least 60% for CASP 13 and CAMEO. At the same time, when the number of retrieved sequences is increased to 2M, fastMSA can successfully detect at least 80% of ground truth for all the three datasets. Furthermore, by comparing with the running time of JackHMMER, we can observe that the retrieval time cost is almost negligible, which clearly demonstrates the benefits of the proposed fastMSA method. Please refer to section 4.3 for more details.

Finally, it is worth pointing out that the proposed model is trained by only using 33k MSAs, which is because of the limitations in the hardware environment, namely computation resources. Even though, the proposed method can still achieve very good performance, as mentioned in the above paragraph. In the future, when more computation resources and advanced GPU machines become available, we will be able to scale up the training scale to millions of MSAs, which could potentially further boost the results by a large margin.

### 4.3 Running Time

In this section, we discuss the effectiveness of our proposed fastMSA method in decreasing the running time that is needed for building MSAs. Fig. 4 provides the results of comparing the running time of directly using vanilla JackHMMER and that of after using our proposed fastMSA method, which clearly demonstrates that the proposed method can effectively accelerate the running speed for various top-K settings. In this figure, the bar chart together with the left axis demonstrates the time needed for building MSAs for the whole CASP14 dataset before and after adopting our method. Correspondingly, the line chart and the right axis directly illustrate the acceleration effect, which indicates how many times that our method can decrease the needed time compared with the original method.

**Figure 4:**
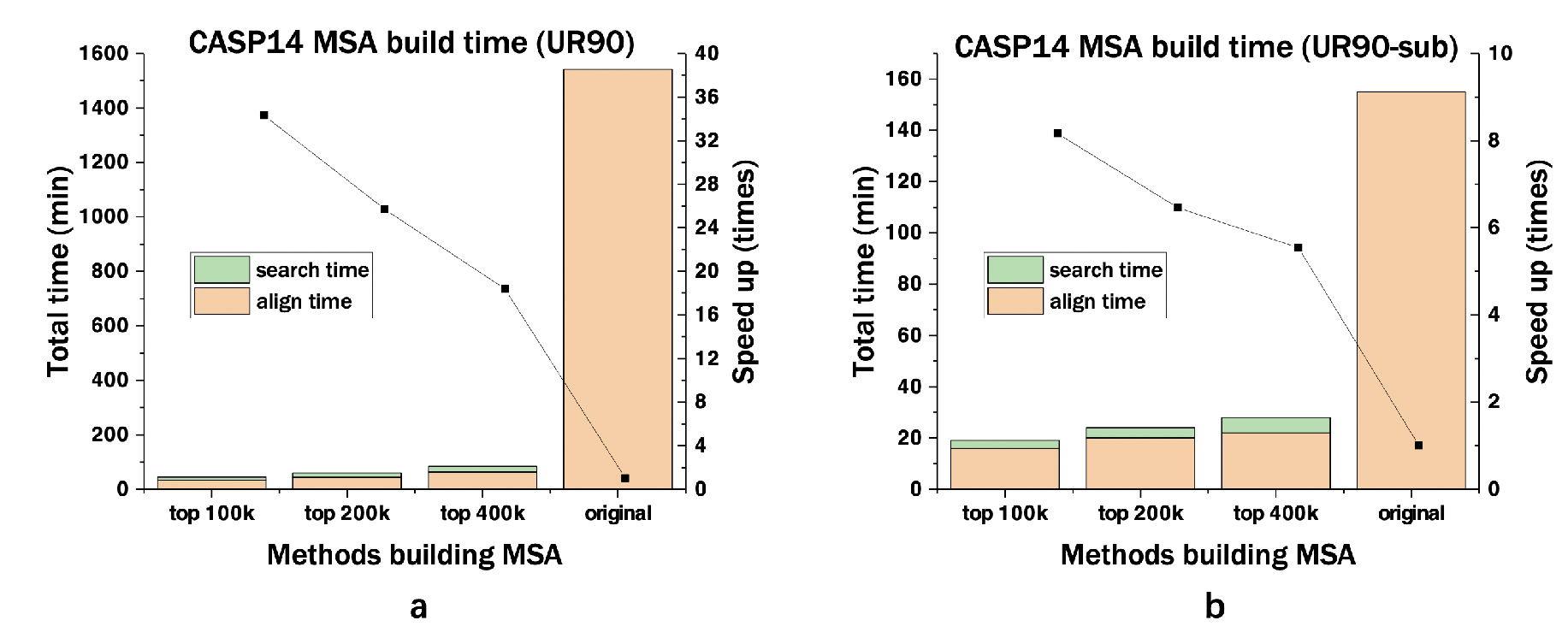
The time taken to build MSAs using JackHMMER. Sub-figure **a** and **b** show the performance on Uniref90 version 2018.3 and the sub-set built by us.

We can clearly observe from this figure that the proposed method can effectively speed up the running speed. For example, for the top 100k sequences, the proposed method achieves a 34 times speedup using our predicted sequences on Uniref90 with around 70M sequences. Since the scanning and the aligning are two major components of JackHMMER searching time, our method can successfully cut the scanning time to minimum. More importantly, a further speedup is expected when the proposed method is utilized to deal with a larger sequence pool.

We also conduct experiences on databases ranging from 10M sequences to 70M and logged the time taken retrieving from 10k to 1M targets shown in Fig. 5 which clearly shows the recall rate grows linearly through time on each database and the time taken in retrieval grows logarithmically with target number.

**Figure 5:**
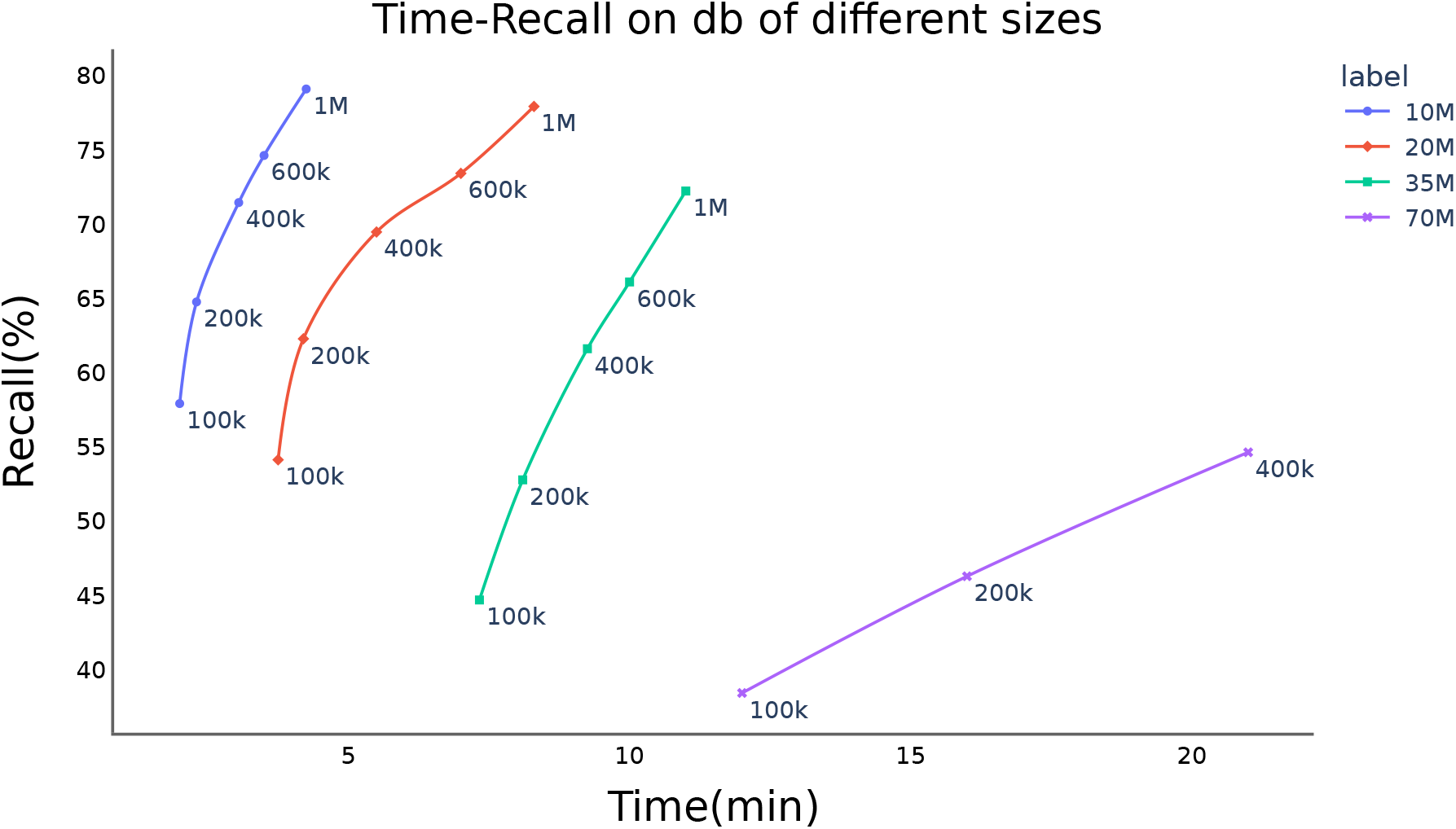
The recall against time taken searching on databases of different sizes and different recall sequences. The legend denotes the size of sequence database and the annotation of point means the retrieve number.

### 4.4 Evaluation on Downstream Tasks

To evaluate the quality of the retrieved MSAs by the proposed method, the 3D structures are built via AlphaFold2 and RoseTTAFold. Instead of using the MSA searching protocols adopted in AlphaFold2 and RoseTTAFold, the MSAs retrieved by our methods are fed into the 3D modeling steps. By default, five 3D models are generated by AlphaFold2, whereas only one model is provided by RoseTTAFold. To evaluate the modeling qualities of the 3D models, the global lDDT score between the native structures and the modeled 3D structures is calculated accordingly.

We present the structure prediction results using retrieved MSA in Table 2. In particular, we compare the AlphaFold2 and RoseTTAFold modeling results that are using (1) original MSA from the whole UniRef90 (original); (2) MSA generated from a subset of UniRef90 that created by ourselves, which serves as the upper bound of the proposed method (ground truth) and (3) MSA generated by proposed method using top 200k and top 1M sequences (top 200k and top 1M). It can be clearly seen from the table that our predicted results has a slight drop compared with the ground truth column predicted by AlphaFold2, but is still comparative to (see results on CAMEO) or even superior to (see results of CASP13 and CASP14) RoseTTAFold results with raw MSA searched against UniRef90. All these indicate the high quality of the MSAs retrieved by the proposed method.

**Table 2:** Modeling precision with corresponding filtered ground truth MSA over 10 sequences

| Dataset | AlphaFold2 |  |  |  | RoseTTAFold |  |
| --- | --- | --- | --- | --- | --- | --- |
|  | original | ground truth | top 200k | top 1M | original | ground truth |
| **CASP13** | 84.69 | 78.90 | 74.47 | **75.49** | 71.83 | 69.12 |
| **CASP14** | 77.28 | 71.20 | **65.86** | 64.14 | 63.91 | 61.78 |
| **CAMEO** | 79.12 | 73.15 | 68.69 | 69.23 | **70.15** | 68.25 |

Finally, we would like to mention an interesting phenomenon that we observe. In our analysis, we find that the modeling results are very sensitive to the build-up of MSAs. Although the top 200k sequences retrieved by our method is only a small subset of the top 1M sequences, the resulted MSA searched from the top 200k sequences could still achieve better performances occasionally, or even for the whole CASP14 dataset. Therefore high recall does not necessarily lead to higher folding accuracy. It demonstrates that our method is able to make a good balance between computation time and the quality of the retrieved MSAs. More importantly, for some datasets, it can further decrease the running time while improving the retrieval quality. We further check this interesting phenomenon in Section 4.7 on three representative proteins by comparing MSAs from our method to the ground-truth MSAs.

### 4.5 Accuracy with respects to Meff

Meff is used since it is a metric for the number of non-redundant sequence homologs in a MSA. Following [47], we use 70% sequence identity as the cutoff to compute the similarity matrix between any two sequences. Please refer to [48, 47] for more detailed formulas. Intuitively, MSAs with small Meff are more difficult to predict since they contain less homologous information.

To check the prediction accuracy using proposed fastMSA with respect to the number of homologs, we further divide MSAs into 7 bins based on their ln(Meff), and compute the average lDDT of ground truth and fastly retrieved MSAs (top 200k and top 1M) for each bin. The results are presented in Fig. 3 b to d. First of all, we notice that the average lDDT increases as ln(Meff) is getting larger from 0 to 5, then flattened after its value is larger than 5 for all three approaches, which might be due to the fact that AlphaFold 2 does not benefit much from more sequence information once it surpasses a threshold, which is consistent with the findings in [22]. In addition, retrieving more sequences to build MSA does not necessarily lead to higher accuracy since the performance on top 200k and top 1M are comparable on almost all bins. This is an interesting topic, and we will explore more in future work. Finally, the lDDT on ground truth MSAs outperforms the proposed fastMSA significantly when ln(Meff) is small in most cases, which indicates that retrieving correct homologs is difficult for such queries.

### 4.6 Head-to-head comparison on CASP13 and CASP14

To evaluate fastMSA more comprehensively, we perform a head-to-head between MSAs from fastMSA and the ground truth MSAs on the protein 3D structure modeling. We use AlphaFold2 to assist the comparison, predicting the 3D structures with it using different MSAs. We evaluate the 3D structure prediction precision for each protein using the ground-truth MSA and the MSA from our method, respectively. When running our method, we filter out the top-200K sequences before performing the standard searching and alignment. We visualize all the results for CASP13 and CASP14 in Fig. 6, with each point representing a protein. The X-axis shows the performance of the ground-truth MSAs, while the Y-axis shows the prediction precision using our MSAs. If the point is far below the red diagonal line, the ground-truth MSA is better for the protein. As shown in the figure, most of the points are around the line, which suggests that our method’s acceleration does not compromise the prediction performance too much. Surprisingly, not all the points are below the red line. Sometimes, our method can even achieve a better prediction precision than the ground-truth MSAs on both the CASP13 and CASP14. In addition to providing acceleration, our method may be combined with traditional MSA methods to further boost the protein structure prediction performance.

**Figure 6:**
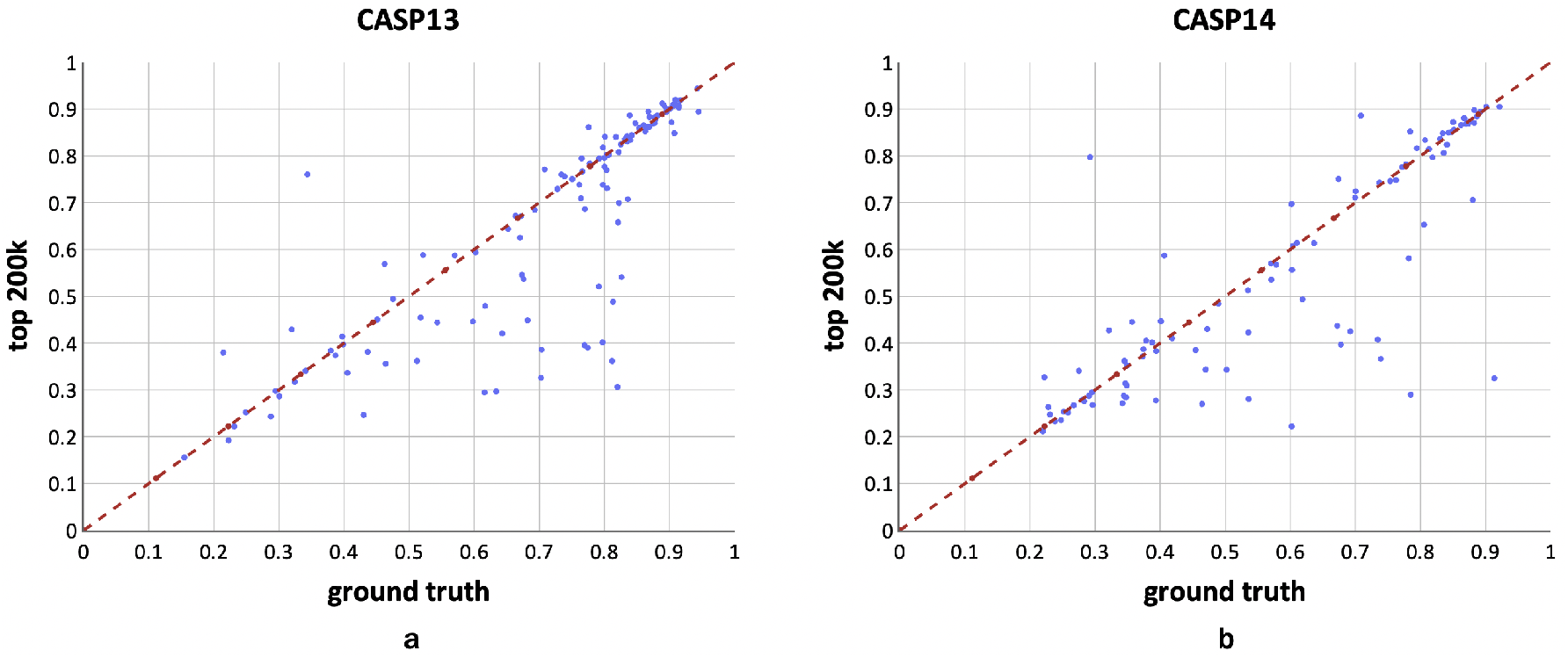
Comparison of 3D structural modeling precision for each protein using the ground-truth MSAs and MSAs from our method. We use AlphaFold2 to predict the 3D structures from the MSAs. **a** Head-to-head comparison between the ground-truth MSAs and MSAs from fastMSA with top-200k filtering on the 3D structural modeling precision for the CASP13 proteins. **b** Head-to-head comparison between the ground-truth MSAs and MSAs from fastMSA with top-200k filtering on the 3D structural modeling precision for the CASP14 proteins.

### 4.7 Case Study

As shown in Fig. 6, the MSAs from fastMSA are comparable to, sometimes even better than, the groundtruth MSAs in 3D structure modeling. We investigate the performance of fastMSA in more detail on three representative proteins from CASP14 dataset by visualizing the predicted 3D structures directly. The structures of three representative targets from the CASP14 dataset are presented in Fig. 7. For both ground-truth MSAs and top 200k MSAs, AF2 successfully builds the high accurate 3D structure models for T1070-D2, with global lDDT scores of 92.13 and 90.49. For T1070-D2, the 3D models from both MSAs are in perfect alignment in not only beta-sheets but also loops regions. For target T1052-D3, the 3D model based on ground-truth MSA is more close to the native structure (global lDDT 91.33), whereas the 3D model is based on top 200k MSA (with a low global lDDT score 32.48) however forms missing orientated *β*-sheets. For the 3rd case, the AF2 3D model based on 200k MSA is more similar to the native structure (global lDDT 79.72) than the 3D model based on the ground-truth MSA (global lDDT 29.24).

**Figure 7:**
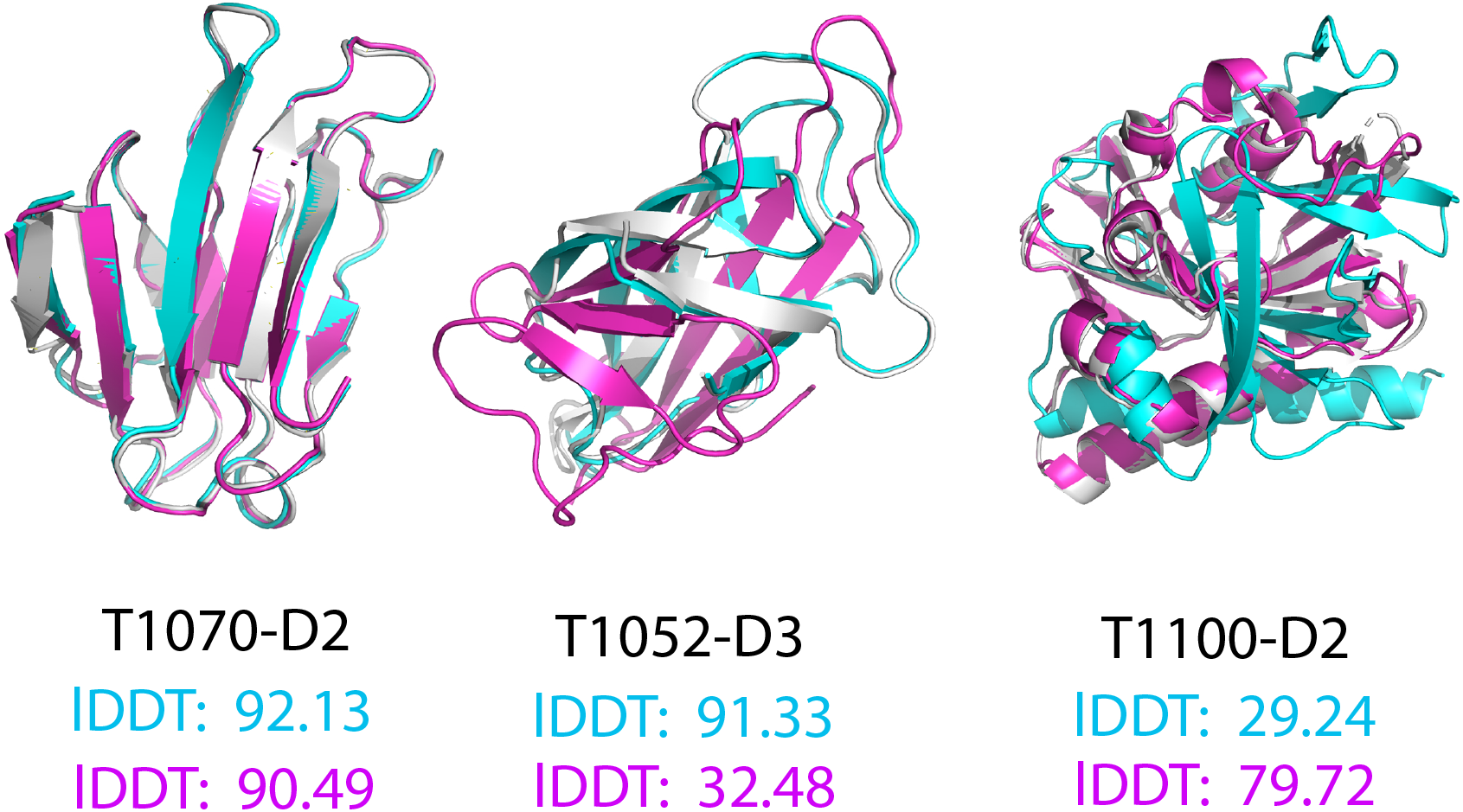
The representative AF2 3D structures based on ground-truth MSAs and top 200k MSAs. The native structures are gray, structures based on ground-truth MSAs are cyan, and structures based on top 200k MSAs are magenta. The values in this figure are global lDDT scores.

## 5 Discussion and Conclusion

MSAs are vital for protein structural and functional analysis. In the post-AlphaFold era, large-scale multiple sequence alignments against massive datasets will be foreseeably popular [49], while its speed is the bottleneck. This paper proposes a new retrieval framework, fastMSA, based on Transformer and contrastive loss to accelerate the MSA generation and its inference time, which is orthogonal to all the previous accelerating methods. By filtering out the unrelated sequences on the low-dimensional space before performing MSA, our method can accelerate the process by 35 folds. As our method is orthogonal to all the other traditional bioinformatic algorithms, fastMAS can be combined with the other methods seamlessly without changing the previous tools and packages. We validate its effectiveness and efficiency on multiple benchmarks and observe promising results and substantial speed improvement on protein structure prediction. Essentially, we can speed up the protein structure prediction using AlphaFold with little performance compromise by accelerating the MSA generation step. In the future, we will expand the size of the MSA training set to obtain more efficient sequence embedding and improve the recall rate further, and apply the proposed method to other downstream tasks, such as protein-RNA interaction modeling [49].

## Notes

### Competing Interest Statement

The authors have declared no competing interest.

## References

[1] Alley, E.C., Khimulya, G., Biswas, S., AlQuraishi, M., Church, G.M.: Unified rational protein engineering with sequence-only deep representation learning. bioRxiv p. 589333 (2019)

[2] Altschul, S.F., Madden, T.L., Schäffer, A.A., Zhang, J., Zhang, Z., Miller, W., Lipman, D.J.: Gapped blast and psi-blast: a new generation of protein database search programs. Nucleic acids research 25(17), 3389–3402 (1997)

[3] Amorim, A.R., Zafalon, G.F.D., de Godoi Contessoto, A., Valêncio, C.R., Sato, L.M.: Metaheuristics for multiple sequence alignment: a systematic review. Computational biology and chemistry p. 107563 (2021)

[4] Armougom, F., Moretti, S., Poirot, O., Audic, S., Dumas, P., Schaeli, B., Keduas, V., Notredame, C.: Expresso: automatic incorporation of structural information in multiple sequence alignments using 3d-coffee. Nucleic acids research 34(suppl 2), W604–W608 (2006)

[5] Bepler, T., Berger, B.: Learning protein sequence embeddings using information from structure. arXiv preprint 1902.08661 (2019)

[6] Bouatta, N., Sorger, P., AlQuraishi, M.: Protein structure prediction by alphafold2: are attention and symmetries all you need? Acta Crystallographica Section D: Structural Biology 77(8), 982–991 (2021)

[7] Buchfink, B., Xie, C., Huson, D.H.: Fast and sensitive protein alignment using diamond. Nature methods 12(1), 59–60 (2015)

[8] Camacho, C., Coulouris, G., Avagyan, V., Ma, N., Papadopoulos, J., Bealer, K., Madden, T.L.: Blast+: architecture and applications. BMC bioinformatics 10(1), 1–9 (2009)

[9] Cao, Y., Geddes, T.A., Yang, J.Y.H., Yang, P.: Ensemble deep learning in bioinformatics. Nature Machine Intelligence 2(9), 500–508 (2020)

[10] Chatzou, M., Magis, C., Chang, J.M., Kemena, C., Bussotti, G., Erb, I., Notredame, C.: Multiple sequence alignment modeling: methods and applications. Briefings in bioinformatics 17(6), 1009–1023 (2016)

[11] Da Costa, M., Gevaert, O., Van Overtveldt, S., Lange, J., Joosten, H.J., Desmet, T., Beerens, K.: Structure-function relationships in ndp-sugar active sdr enzymes: Fingerprints for functional annotation and enzyme engineering. Biotechnology Advances p. 107705 (2021)

[12] Daugelaite, J., O’Driscoll, A., Sleator, R.D.: An overview of multiple sequence alignments and cloud computing in bioinformatics. International Scholarly Research Notices 2013 (2013)

[13] Devlin, J., Chang, M.W., Lee, K., Toutanova, K.: Bert: Pre-training of deep bidirectional transformers for language understanding. arXiv preprint 1810.04805 (2018)

[14] Do, C.B., Mahabhashyam, M.S., Brudno, M., Batzoglou, S.: Probcons: Probabilistic consistency-based multiple sequence alignment. Genome research 15(2), 330–340 (2005)

[15] Edgar, R.C.: Muscle: multiple sequence alignment with high accuracy and high throughput. Nucleic acids research 32(5), 1792–1797 (2004)

[16] Edgar, R.C., Batzoglou, S.: Multiple sequence alignment. Current opinion in structural biology 16(3), 368–373 (2006)

[17] Fukuda, H., Tomii, K.: Deepeca: an end-to-end learning framework for protein contact prediction from a multiple sequence alignment. BMC bioinformatics 21(1), 1–15 (2020)

[18] Heinzinger, M., Elnaggar, A., Wang, Y., Dallago, C., Nechaev, D., Matthes, F., Rost, B.: Modeling the language of life–deep learning protein sequences. bioRxiv p. 614313 (2019)

[19] Hogeweg, P., Hesper, B.: The alignment of sets of sequences and the construction of phyletic trees: an integrated method. Journal of molecular evolution 20(2), 175–186 (1984)

[20] Huson, D.H., Tappu, R., Bazinet, A.L., Xie, C., Cummings, M.P., Nieselt, K., Williams, R.: Fast and simple protein-alignment-guided assembly of orthologous gene families from microbiome sequencing reads. Microbiome 5(1), 1–10 (2017)

[21] Johnson, J., Douze, M., Jégou, H.: Billion-scale similarity search with gpus. IEEE Transactions on Big Data (2019)

[22] Jumper, J., Evans, R., Pritzel, A., Green, T., Figurnov, M., Ronneberger, O., Tunyasuvunakool, K., Bates, R., Ž ídek, A., Potapenko, A., et al.: Highly accurate protein structure prediction with alphafold. Nature 596(7873), 583–589 (2021)

[23] Karpukhin, V., Oğuz, B., Min, S., Lewis, P., Wu, L., Edunov, S., Chen, D., Yih, W.t.: Dense passage retrieval for open-domain question answering. arXiv preprint 2004.04906 (2020)

[24] Katoh, K., Misawa, K., Kuma, K.i., Miyata, T.: Mafft: a novel method for rapid multiple sequence alignment based on fast fourier transform. Nucleic acids research 30(14), 3059–3066 (2002)

[25] Kececioglu, J.D., Lenhof, H.P., Mehlhorn, K., Mutzel, P., Reinert, K., Vingron, M.: A polyhedral approach to sequence alignment problems. Discrete applied mathematics 104(1-3), 143–186 (2000)

[26] Lesk, A.M., Chothia, C.: How different amino acid sequences determine similar protein structures: the structure and evolutionary dynamics of the globins. Journal of molecular biology 136(3), 225–270 (1980)

[27] Lewis, P., Perez, E., Piktus, A., Petroni, F., Karpukhin, V., Goyal, N., Küttler, H., Lewis, M., Yih, W.t., Rocktäschel, T., et al.: Retrieval-augmented generation for knowledge-intensive nlp tasks. arXiv preprint 2005.11401 (2020)

[28] Li, Y., Huang, C., Ding, L., Li, Z., Pan, Y., Gao, X.: Deep learning in bioinformatics: Introduction, application, and perspective in the big data era. Methods 166, 4–21 (2019)

[29] Li, Y., Wang, S., Umarov, R., Xie, B., Fan, M., Li, L., Gao, X.: Deepre: sequence-based enzyme ec number prediction by deep learning. Bioinformatics 34(5), 760–769 (2018)

[30] Makigaki, S., Ishida, T.: Sequence alignment using machine learning for accurate template-based protein structure prediction. Bioinformatics 36(1), 104–111 (2020)

[31] Murtagh, F.: Complexities of hierarchic clustering algorithms: state of the art. Computational Statistics Quarterly 1(2), 101–113 (1984)

[32] Needleman, S.B., Wunsch, C.D.: A general method applicable to the search for similarities in the amino acid sequence of two proteins. Journal of molecular biology 48(3), 443–453 (1970)

[33] Notredame, C., Higgins, D.G., Heringa, J.: T-coffee: A novel method for fast and accurate multiple sequence alignment. Journal of molecular biology 302(1), 205–217 (2000)

[34] O’Sullivan, O., Suhre, K., Abergel, C., Higgins, D.G., Notredame, C.: 3dcoffee: combining protein sequences and structures within multiple sequence alignments. Journal of molecular biology 340(2), 385–395 (2004)

[35] Radford, A., Wu, J., Child, R., Luan, D., Amodei, D., Sutskever, I., et al.: Language models are unsupervised multitask learners. OpenAI blog 1(8), 9 (2019)

[36] Rao, R., Meier, J., Sercu, T., Ovchinnikov, S., Rives, A.: Transformer protein language models are unsupervised structure learners. In: International Conference on Learning Representations (2020)

[37] Rives, A., Meier, J., Sercu, T., Goyal, S., Lin, Z., Liu, J., Guo, D., Ott, M., Zitnick, C.L., Ma, J., et al.: Biological structure and function emerge from scaling unsupervised learning to 250 million protein sequences. Proceedings of the National Academy of Sciences 118(15) (2021)

[38] Saitou, N., Nei, M.: The neighbor-joining method: a new method for reconstructing phylogenetic trees. Molecular biology and evolution 4(4), 406–425 (1987)

[39] Senior, A.W., Evans, R., Jumper, J., Kirkpatrick, J., Sifre, L., Green, T., Qin, C., Zídek, A., Nelson, A.W.R., Bridgland, A., Penedones, H., Petersen, S., Simonyan, K., Crossan, S., Kohli, P., Jones, D.T., Silver, D., Kavukcuoglu, K., Hassabis, D.: Improved protein structure prediction using potentials from deep learning - Nature. Nature 577, 706–710 (2020)

[40] Sievers, F., Wilm, A., Dineen, D., Gibson, T.J., Karplus, K., Li, W., Lopez, R., McWilliam, H., Remmert, M., et al.: Fast, scalable generation of high-quality protein multiple sequence alignments using clustal omega. Molecular systems biology 7(1), 539 (2011)

[41] Steinegger, M., Meier, M., Mirdita, M., Vöhringer, H., Haunsberger, S.J., Söding, J.: Hh-suite3 for fast remote homology detection and deep protein annotation. BMC bioinformatics 20(1), 1–15 (2019)

[42] Thompson, J.D., Higgins, D.G., Gibson, T.J.: Clustal w: improving the sensitivity of progressive multiple sequence alignment through sequence weighting, position-specific gap penalties and weight matrix choice. Nucleic acids research 22(22), 4673–4680 (1994)

[43] Thompson, J.D., Linard, B., Lecompte, O., Poch, O.: A comprehensive benchmark study of multiple sequence alignment methods: current challenges and future perspectives. PloS one 6(3), e18093 (2011)

[44] Thornton, J.M., Laskowski, R.A., Borkakoti, N.: Alphafold heralds a data-driven revolution in biology and medicine. Nature Medicine pp. 1–4 (2021)

[45] Wallace, I.M., Higgins, D.G.: Evaluation of iterative alignment algorithms for multiple alignment. Bioinformatics 21(8), 1408–1414 (2005)

[46] Wang, S., Fei, S., Wang, Z., Li, Y., Xu, J., Zhao, F., Gao, X.: Predmp: a web server for de novo prediction and visualization of membrane proteins. Bioinformatics 35(4), 691–693 (2019)

[47] Wang, S., Sun, S., Li, Z., Zhang, R., Xu, J.: Accurate de novo prediction of protein contact map by ultra-deep learning model. PLoS computational biology 13(1), e1005324 (2017)

[48] Wang, Z., Xu, J.: Predicting protein contact map using evolutionary and physical constraints by integer programming. Bioinformatics 29(13), i266–i273 (Jul 2013). https://doi.org/10.1093/bioinformatics/btt211

[49] Wei, J., Chen, S., Zong, L., Gao, X., Li, Y.: Protein-rna interaction prediction with deep learning: Structure matters. arXiv preprint 2107.12243 (2021)

[50] Wheeler, T.J., Eddy, S.R.: nhmmer: Dna homology search with profile hmms. Bioinformatics 29(19), 2487–2489 (2013)

[51] Xia, X., Zhang, S., Su, Y., Sun, Z.: Micalign: a sequence-to-structure alignment tool integrating multiple sources of information in conditional random fields. Bioinformatics 25(11), 1433–1434 (2009)

[52] Xiong, L., Xiong, C., Li, Y., Tang, K.F., Liu, J., Bennett, P., Ahmed, J., Overwijk, A.: Approximate nearest neighbor negative contrastive learning for dense text retrieval. arXiv preprint 2007.00808 (2020)

[53] Yazdani, Z., Rafiei, A., Valadan, R., Ashrafi, H., Pasandi, M., Kardan, M.: Designing a potent l1 protein-based hpv peptide vaccine: a bioinformatics approach. Computational biology and chemistry 85, 107209 (2020)

[54] Zhang, Y., Sun, S., Gao, X., Fang, Y., Brockett, C., Galley, M., Gao, J., Dolan, B.: Joint retrieval and generation training for grounded text generation. arXiv preprint 2105.06597 (2021)

[55] Zou, Q., Lin, G., Jiang, X., Liu, X., Zeng, X.: Sequence clustering in bioinformatics: an empirical study. Briefings in bioinformatics 21(1), 1–10 (2020)

[56] Zou, Z., Tian, S., Gao, X., Li, Y.: mldeepre: Multi-functional enzyme function prediction with hierarchical multi-label deep learning. Frontiers in Genetics 9, 714 (2019)

